# Heteologous saRNA-Prime, DNA Dual-Antigen-Boost SARS-CoV-2 Vaccination Elicits Robust Cellular Immunogenicity and Cross-Variant Neutralizing Antibodies

**DOI:** 10.1101/2021.11.29.470440

**Authors:** Adrian Rice, Mohit Verma, Emily Voigt, Peter Battisti, Sam Beaver, Sierra Reed, Kyle Dinkins, Shivani Mody, Lise Zakin, Shiho Tanaka, Brett Morimoto, C. Anders Olson, Elizabeth Gabitzsch, Jeffrey T. Safrit, Patricia Spilman, Corey Casper, Patrick Soon-Shiong

**Author notes:** Corresponding author Patrick Soon-Shiong. These authors contributed equally (Adrian Rice, Mohit Verma, Emily Voigt).

## Abstract

We assessed if immune responses are enhanced in CD-1 mice by heterologous vaccination with two different nucleic acid-based COVID-19 vaccines: a next-generation human adenovirus serotype 5 (hAd5)-vectored dual-antigen spike (S) and nucleocapsid (N) vaccine (AdS+N) and a self-amplifying and -adjuvanted S RNA vaccine (SASA S) delivered by a nano-lipid carrier. The AdS+N vaccine encodes S modified with a fusion motif to increase cell-surface expression. The N antigen is modified with an Enhanced T-cell Stimulation Domain (N-ETSD) to direct N to the endosomal/lysosomal compartment and increase MHC class I and II stimulation potential. The S sequence in the SASA S vaccine comprises the D614G mutation, two prolines to stabilize S in the prefusion conformation, and 3 glutamines in the furin cleavage region to increase cross-reactivity across variants. CD-1 mice received vaccination by homologous and heterologous prime > boost combinations. Humoral responses to S were the highest with any regimen including the SASA S vaccine, and IgG bound to wild type and Delta (B.1.617.2) variant S1 at similar levels. An AdS+N boost of an SASA S prime particularly enhanced both CD4+ and CD8+ T-cell responses to both wild type and Delta S peptides relative to all other vaccine regimens. Sera from mice receiving SASA S homologous or heterologous vaccination were found to be highly neutralizing of all pseudovirus strains tested: Wuhan, Beta, Delta, and Omicron strain. The findings here support the clinical testing of heterologous vaccination by an SASA S > AdS+N regimen to provide increased protection against emerging SARS-CoV-2 variants.

## INTRODUCTION

Impressive efforts of the scientific and pharmaceutical community have resulted in the design, testing and successful deployment of several COVID-19 vaccines that have shown high levels of efficacy [1–5]. Nonetheless, SARS-CoV-2 viral variants have continued to emerge and spread throughout the globe – most recently the highly transmissible Omicron variant [6] – pointing to the need for delivery of vaccines to populations that are currently underserved.

To address the need for a vaccine regimen that would be highly efficacious against predominating and emerging variants as well as distributable in currently underserved areas, we previously developed a next-generation human adenovirus serotype 5 (hAd5)-vectored dual-antigen spike (S) plus nucleocapsid (N) vaccine (AdS+N) [7, 8] to leverage the resilience of cell-mediated immunity against variants. This vaccine, encoding Wuhan strain or ‘wild type’ (wt) SARS-CoV-2 S and modified with a fusion sequence (S-Fusion) to enhance cell-surface expression [7, 8], as well as N modified with an Enhanced T-cell Stimulation Domain (N-ETSD) [9] for increased MHC class I and II stimulation [10–12], has been shown to elicit humoral and T-cell responses in mice, [8] non-human primates (NHP) [7], and participants in Phase 1b trials [9]. The Ad5S+N vaccine given as a subcutaneous (SC) prime with two oral boosts protected NHP from SARS-CoV-2 infection [7], and a single prime vaccination of clinical trial participants generated T-cell responses that were sustained against a series of variant S peptide sequences, including those for the B.1.351, B.1.1.7, P.1, and B.1.426 variants [9].

Despite the promising findings with the AdS+N vaccine candidate, we wish to continue to investigate vaccine regimens with the potential to maximize immune responses – both humoral and cellular. One such approach is by heterologous vaccination utilizing multiple nucleic acid-based vaccine platforms, such as ImmunityBio’s hAd5 vectored DNA vaccine and the Infectious Disease Research Institute’s (IDRI’s) RNA-based vaccine [13]. Heterologous vaccination using vaccine constructs expressing the same or different antigens vectored by different platforms, specifically combinations of RNA- and adenovirus-based vaccines, has previously been reported to significantly increase immune responses [14, 15].

To assess the potential for enhanced immune responses by heterologous vaccination, we tested prime > boost combinations of the AdS+N vaccine with a self-amplifying and self-adjuvanted S(wt) RNA-based vaccine (SASA S) delivered in a nanolipid carrier (NLC) [16, 17]. The NLC stabilizes the self-amplifying RNA [18–20] and delivers it to cells, where the vaccine RNA is then amplified and S protein expressed. The S sequence in the SASA S vaccine comprises a codon-optimized sequence with the D614G mutation [21] that increases SARS-CoV-2 susceptibility to neutralization [22], a diproline modification to stabilize S in the pre-fusion conformation that increases antigenicity [23], and a tri-glutamine (3Q) repeat in the furin cleavage region that appears to broaden immune responses against SARS-CoV-2 variants [24]. Preclinical studies of the SASA S vaccine have demonstrated the vaccine elicits vigorous antigen-specific and virus-neutralizing IgG and polyfunctional CD4+ and CD8+ T-cell responses after both prime and prime-boost regimens in C57Bl/6 mice.

In this work, the two aforementioned vaccines were tested by homologous and heterologous AdS+N > SASA S and SASA S > AdS+N prime > boost regimens:. The findings reported here support our hypothesis that heterologous vaccination with the SASA S and AdS+N vaccines enhances immune responses, and particularly T-cell responses.

## METHODS

### The AdS+N and SASA S vaccines

For studies here, the next generation hAd5 [E1 -, E2b-, E3-] vector was used to create viral vaccine candidate constructs [7]. This hAd5 [E1-, E2b-, E3-] vector is primarily distinguished from other first-generation [E1-, E3-] recombinant Ad5 platforms [25, 26] by having additional deletions in the early gene 2b (E2b) region that remove the expression of the viral DNA polymerase (pol) and in pre terminal protein (pTP) genes, and its propagation in the E.C7 human cell line [27–30].

The AdS+N vaccine expresses a wild type spike (S) sequence [accession number YP009724390] modified with a proprietary ‘fusion’ linker peptide sequence as well as a wild type nucleocapsid (N) sequence [accession number YP009724397] with an Enhanced T-cell Stimulation Domain (ETSD) signal sequence to direct translated N to the endosomal/lysosomal pathway [9] as described in Gabitzsch *et al.,* 2021 [7].

The SASA S vaccine comprises an saRNA replicon composed of an 11.7 kb construct expressing the SARS-CoV-2 Spike protein, along with the non-structural proteins 1-4 derived from the Venezuelan equine encephalitis virus (VEEV) vaccine strain TC-83. The Spike RNA sequence is codon-optimized and expresses a protein with the native sequence of the original Wuhan strain plus the dominant D614G mutation, with the prefusion conformation-stabilizing diproline (pp) mutation (consistent with other vaccine antigens) and replacement of the furin cleavage site RRAR sequence with a QQAQ sequence, as shown in **Figure 1**.

**Fig. 1.**
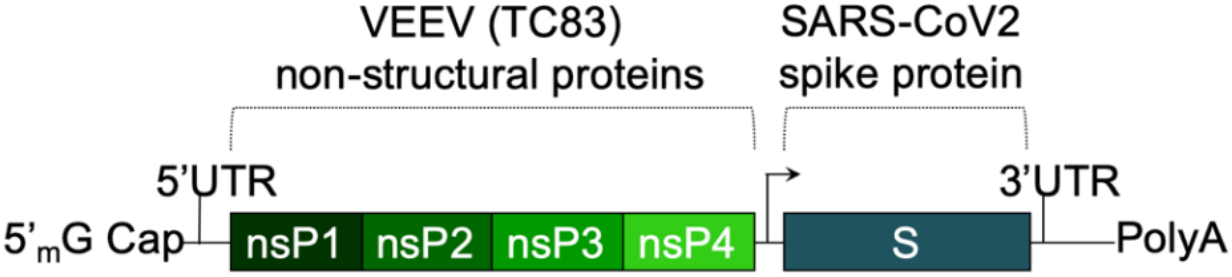
The saRNA(D614G)-2P-3Q-NLC (SASA S) vaccine. The SASA S vaccine comprises an saRNA replicon backbone consisting of the non-structural protein (nsPs) 1-4 derived from the Venezuelan equine encephalitis virus (VEEV) vaccine strain TC-83 and an independent open reading frame under the control of a subgenomic promoter sequence that contains Wuhan sequence S with a diproline (pp) mutation and a QQAQ furin cleavage site sequence.

The RNA is generated by T7 promoter-mediated *in vitro* transcription using a linearized DNA template. *In vitro* transcription is performed using an in house-optimized protocol [13, 31, 32] using T7 polymerase, RNase inhibitor, and pyrophosphatase enzymes. The DNA plasmid is digested with DNase I and the RNA is capped by vaccinia capping enzyme, guanosine triphosphate, and S-adenosyl-methionine. RNA is then purified from the transcription and capping reaction components by chromatography using a CaptoCore 700 resin (GE Healthcare) followed by diafiltration and concentration using tangential flow filtration into 10 mM Tris buffer. The RNA material is terminally filtered with a 0.22 μm polyethersulfone filter and stored at −80°C until use.

The RNA-delivering NLC is comprised of particles with a hybrid liquid and solid oil core, providing colloidal stability [33], surrounded by non-ionic hydrophobic and hydrophilic surfactants to help maintain a stable nanoparticle droplet and the cationic lipid DOTAP to provide positive charge for electrostatic binding with RNA. This RNA binding on the surface of the nanoparticles protects the RNA RNase degradation and allows effective delivery to cells.

NLC is manufactured by mixing the lipids in an oil phase, dissolving the Tween 80 in citrate buffer aqueous phase, and homogenizing the two phases by micro-fluidization. The resulting emulsion is sterile-filtered and vialed until dilution in a sucrose-citrate solution and complexing with vaccine saRNA.

### Murine immunization and blood/tissue collection

The design of vaccination study performed using CD-1 mice is shown in **Figure 2**.

**Fig. 2.**
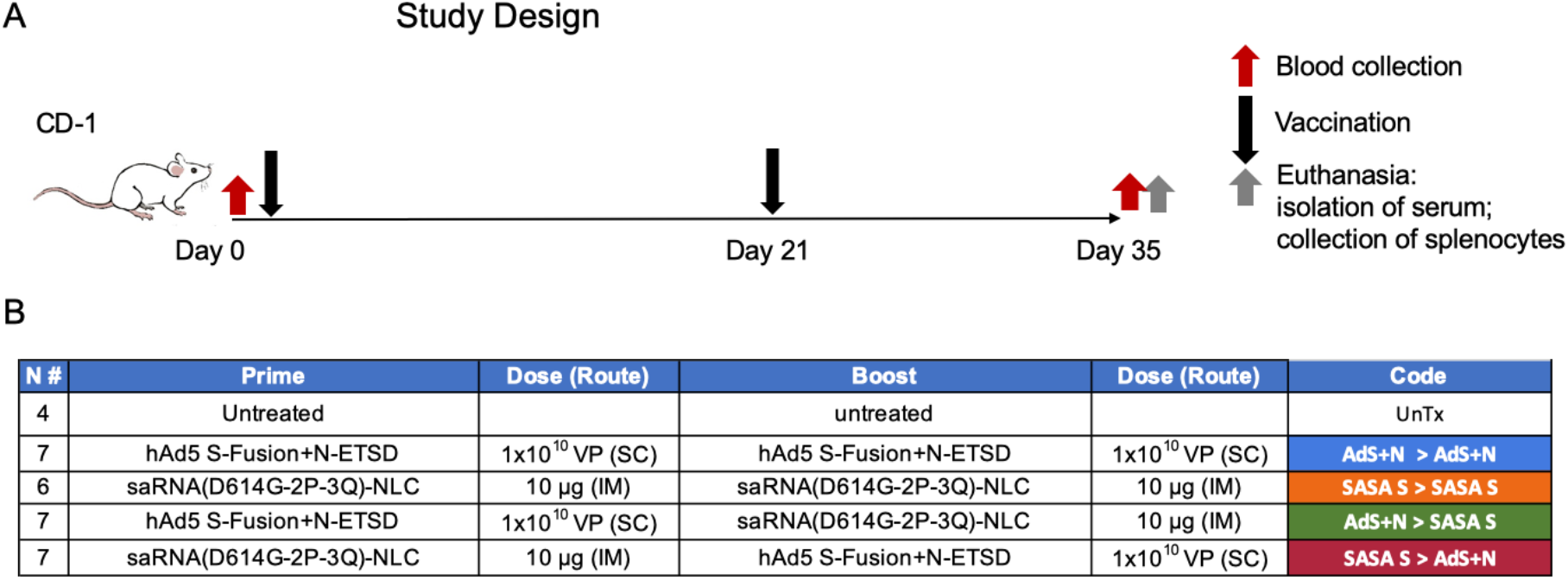
Study design and vaccine description. (A) CD-1 mice received prime vaccination on Day 0 after blood collection and boost vaccination on Day 21; mice were euthanized and tissues/blood collected on Day 35. (B) The various combinations of prime > boost are shown, including: AdS+N homologous; saRNA(D614G-2P-3Q)-NLC (SASA S) homologous; Ad5S+N prime, SASA S boost; and SASA S prime, AdS+N boost. Untreated mice were used as controls. All groups were n = 7 with the exception of untreated n = 4 and SASA S homologous n = 6. The color code for each group is shown.

All *in vivo* experiments described were carried out at the Omerios Inc. vivarium (Seattle, WA) in strict accordance with good animal practice according to NIH recommendations. All procedures for animal use were done under an animal use protocol approved by the IACUC at Omeros, Inc. (Seattle, WA, USA).

CD-1 female mice (Charles River Laboratories) 6-8 weeks of age were used for immunological studies. The adenovirus-vectored vaccines were administered by subcutaneous (SC) injections at the indicated doses in 50 μL ARM buffer (20 mM Tris pH 8.0, 25 mM NaCl, with 2.5% glycerol). The SASA S vaccine was administered intramuscularly (IM) in 10% sucrose 5 mM sodium citrate solution at a dose of 10 μg.

On the final day of each study, blood was collected submandibularly from isoflurane-anesthetized mice and sera isolated using a microtainer tube. Mice were then euthanized for collection of spleens. Spleens were placed in 5 mL of sterile media (RPMI/HEPES/Pen/Strep/10% FBS). Splenocytes were isolated [34] within 2 hours of collection and used fresh or cryopreserved for later analysis.

### Intracellular cytokine stimulation (ICS)

ICS assays were performed using 10^6^ live splenocytes per well in 96-well U-bottom plates. Splenocytes in RPMI media supplemented with 10% FBS were stimulated by the addition of pools of overlapping peptides spanning the SARS-CoV-2 S protein (both wild type Wuhan strain, wt, or Delta sequence) or N antigens at 2 μg/mL/peptide for 6 h at 37°C in 5% CO2, with protein transport inhibitor, GolgiStop (BD) added two hours after initiation of incubation. The S peptide pool (wild type, JPT Cat #PM-WCPV-S-1; Delta, JPT cat# PM-SARS2-SMUT06-1) is a total of 315 spike peptides split into two pools, S1 and S2, comprised of 158 and 157 peptides each. The N peptide pool (JPT; Cat # PM-WCPV-NCAP-1) was also used to stimulate cells. A SIV-Nef peptide pool (BEI Resources) was used as an off-target negative control. Stimulated splenocytes were then stained with a fixable cell viability stain (eBioscience™ Fixable Viability Dye eFluor™ 506 Cat# 65-0866-14) followed by the lymphocyte surface markers CD8β and CD4, fixed with CytoFix (BD), permeabilized, and stained for intracellular accumulation of IFN-γ, TNF-α and IL-2. Fluorescent-conjugated anti-mouse antibodies used for labeling included CD8β antibody (clone H35-17.2, ThermoFisher), CD4 (clone RM4-5, BD), IFN-γ (clone XMG1.2, BD), TNF-α (clone MP6-XT22, BD) and IL-2 (clone JES6-5H4; BD), and staining was performed in the presence of unlabeled anti-CD16/CD32 antibody (clone 2.4G2; BD). Flow cytometry was performed using a Beckman-Coulter Cytoflex S flow cytometer and analyzed using Flowjo Software.

### ELISpot assay

ELISpot assays were used to detect cytokines secreted by splenocytes from inoculated mice. Fresh splenocytes were used on the same day as harvest, and cryopreserved splenocytes containing lymphocytes were used the day of thawing. The cells (2-4 x 10^5^ cells per well of a 96-well plate) were added to the ELISpot plate containing an immobilized primary antibody to either IFN-g or IL-4 (BD Cat# 551881 and BD Cat# 551878, respectively), and were exposed to various stimuli (e.g. control peptides SIV and ConA, S-WT and N peptides pools – see catalog numbers above) at a concentration of 1-2 μg/mL peptide pools for 36-40 hours. After aspiration and washing to remove cells and media, extracellular cytokines were detected by a biotin-conjugated secondary antibody to cytokine conjugated to biotin (BD), followed by a streptavidin/horseradish peroxidase conjugate was used to detect the biotin-conjugated secondary antibody. The number of spots per well, or per 2-4 x 10^5^ cells, was counted using an ELISpot plate reader. Quantification of Th1/Th2 bias was calculated by dividing the IFN-g spot forming cells (SFC) per million splenocytes with the IL-4 SFC per million splenocytes for each animal.

### ELISA for detection of antibodies

For IgG antibody detection in inoculated mouse sera and lung homogenates, ELISAs for spike-binding (including S1 Delta) and nucleocapsid-binding IgG and IgG subclass (IgG1, IgG2a, IgG2b, and IgG3) antibodies) were used. A microtiter plate was coated overnight with 100 ng of either purified recombinant SARS-CoV-2 S-FTD (FL S with fibritin trimerization domain, constructed and purified in-house by ImmunityBio), purified recombinant Spike S1 domain (S1(wt)) (Sino; Cat # 40591-V08B1), purified recombinant Delta variant Spike S1 domain (S1(Delta)) (Sino; Cat # 40591-V08H23), or purified recombinant SARS-CoV-2 nucleocapsid (N) protein (Sino; Cat # 40588-V08B) in 100 μL of coating buffer (0.05 M Carbonate Buffer, pH 9.6). The wells were washed three times with 250 μL PBS containing 1% Tween 20 (PBST) to remove unbound protein and the plate was blocked for 60 minutes at room temperature with 250 μL PBST. After blocking, the wells were washed with PBST, 100 μL of either diluted serum or diluted lung homogenate samples was added to each well, and samples incubated for 60 minutes at room temperature. After incubation, the wells were washed with PBST and 100 μL of a 1/5000 dilution of anti-mouse IgG HRP (GE Health Care; Cat # NA9310V), anti-mouse IgG1 HRP (Sigma; Cat # SAB3701171), anti-mouse IgG2a HRP (Sigma; Cat # SAB3701178), anti-mouse IgG2b HRP (Sigma; catalog# SAB3701185), anti-mouse IgG3 HRP conjugated antibody (Sigma; Cat # SAB3701192), or anti-mouse IgA HRP conjugated antibody (Sigma; Cat # A4789) was added to wells. For positive controls, 100 μL of a 1/5000 dilution of rabbit anti-N IgG Ab or 100 μL of a 1/25 dilution of mouse antiS serum (from mice immunized with purified S antigen in adjuvant) were added to appropriate wells. After incubation at room temperature for 1 hour, the wells were washed with PBS-T and incubated with 200 μL o-phenylenediamine-dihydrochloride (OPD substrate, Thermo Scientific Cat # A34006) until appropriate color development. The color reaction was stopped with addition of 50 μL 10% phosphoric acid solution (Fisher Cat # A260-500) in water and the absorbance at 490 nm was determined using a microplate reader (SoftMax Pro, Molecular Devices).

### Calculation of relative ng amounts of antibodies and the Th1/Th2 IgG subclass bias

A standard curve of IgG for OD vs. ng mouse IgG was generated using purified mouse IgG (Sigma Cat #15381); absorbance values from this standard curve were used to convert sample absorbance signals into mass equivalents for both anti-S and anti-N antibodies. Using these values, we calculated the geometric mean value for S- and N-specific IgG per milliliter of serum induced by vaccination. These values were also used to quantify the Th1/Th2 bias for the humoral responses by dividing the sum total of Th1 biased antigen-specific IgG subclasses (IgG2a, IgG2b and IgG3) with the total Th2 indicative IgG1, for each mouse. For mice that lacked anti-S and/or anti-N specific IgG responses, Th1/Th2 ratio was not calculated. Some responses, particularly for anti-N responses in IgG2a and IgG2b (both Th1 biased subclasses), were above the limit of quantification with OD values higher than those observed in the standard curve. These data points were therefore reduced to values within the standard curve, and thus the reported Th1/Th2 bias is lower than would otherwise be reported.

### Endpoint titers

Serial dilutions were prepared from each serum sample, with dilution factors ranging from 400 to 6,553,600 in 4-fold steps. These dilution series were characterized by whole IgG ELISA assays against both recombinant S1(wt) and recombinant S1(Delta), as described above. Half maximal response values (Ab50) were calculated by non-linear least squares fit analysis on the values for each dilution series against each recombinant S1 in GraphPad Prism. Serum samples from mice without anti-S responses were removed from Ab50, μg IgG/mL sera, and endpoint titer analyses and reported as N/D on the graphs. Endpoint titers were defined as the last dilution with an absorbance value at least 3 standard deviations higher than the standard deviation of all readings from serum of untreated animals (n = 32 total negative samples). Quantitative titration values (μg IgG/mL sera) were calculated against a standard curve as described above.

### Pseudovirus neutralization assay

SARS-CoV-2 pseudovirus neutralization assays were conducted on immunized mouse serum samples using procedures adapted from Crawford *et al*., 2020. [35] In brief, lentiviral pseudoviruses expressing SARS-CoV-2 spike protein variants were prepared by co-transfecting HEK293 cells (ATCC CRL-3216) seeded at 4*10^5^ cells/mL with a plasmid containing a lentiviral backbone expressing luciferase and ZsGreen (BEI Resources NR-52516), plasmids containing lentiviral helper genes (BEI Resources NR-52517, NR-52518, NR-52519), a delta19 cytoplasmic tail-truncated SARS-CoV-2 spike protein expression plasmid (Wuhan strain, B.1.1.7, and B.1.351 spike variant plasmids were a gift from Jesse Bloom of Fred Hutchinson Cancer Research Center; B.1.617.2 “delta” and omicron variant plasmids a gift from Thomas Peacock of Imperial College London) and Bio-T transfection reagent (Bioland Scientific B0101). The transfection was incubated for 72 hours at 37°C, 5% CO2. Pseudovirus stocks were harvested from the cell culture media, (Gibco DMEM + GlutaMAX + 10% FBS) filtered through a 0.2 μm filter, and frozen at −80°C until titering and use.

Mouse serum samples were diluted 1:10 in media (Gibco DMEM + GlutaMAX + 10% FBS) and then serially diluted 1:2 for 11 total dilutions, and incubated for 1 hour at 37°C, 5% CO2.with a mixture of 5 μg/mL polybrene (Sigma TR-1003-G) and pseudovirus diluted to a titer that produces 1*10^8^ total integrated intensity units/mL. The serum-virus mix was then added in duplicate to human Angiotensin-Converting Enzyme 2 expressing HEK293 cells (BEI Resources NR-52511, NIAID, NIH) seeded at 4 x 10^5^ cell/mL on a 96 well plate.

The plates were incubated at 37°C, 5% CO2 for 72 hours, Plates were imaged on a high content fluorescent imager (Molecular Devices ImageXpress Pico) for ZsGreen expression. Total integrated intensity units per well quantified using ImageXpress software (Molecular Devices) was used to calculate % pseudovirus inhibition in each well. Neutralization curves were fit with a four-parameter sigmoidal curve which was used to calculate 50% inhibitory concentration dilution (IC50) values.

### Statistical analyses and graph generation

All statistical analyses were performed and figures and graphs generated using GraphPad Prism software. Statistical tests for each graph are described in the figure legends. Statistical analyses of Endpoint Titers for anti-S1 IgG (**Fig. 4**) were performed by assigning a value of 200 – one half the Level of Detection (LOD) of 400 – to the 4 animals with serum values below the LOD.

## RESULTS

### The SASA S Vaccine Enhanced Generation of Anti-S(wt) IgG

Mice receiving the SASA S vaccine in any homologous or heterologous vaccination regimen had the highest levels of anti-full length S(wt) (FL S) IgG2a and 2b as determined by ELISA OD readouts OD at 490 nm (**Fig. 3A**). As expected, only mice receiving the N antigen generated anti-N IgG (also determined by ELISA 290 nm OD readout), at levels similar for all groups receiving an N-containing antigen (**Fig. 3B**) by AdS+ N homologous, prime, or boost vaccination. Determination of the IgG1/IgG2a + IgG2b + IgG3 ratio using ng amounts calculated from the OD reading (see *Methods*) revealed responses were highly T helper cell 1 (Th1)-biased, with all calculated values being greater than one (**Fig. 3C**).

**Fig. 3.**
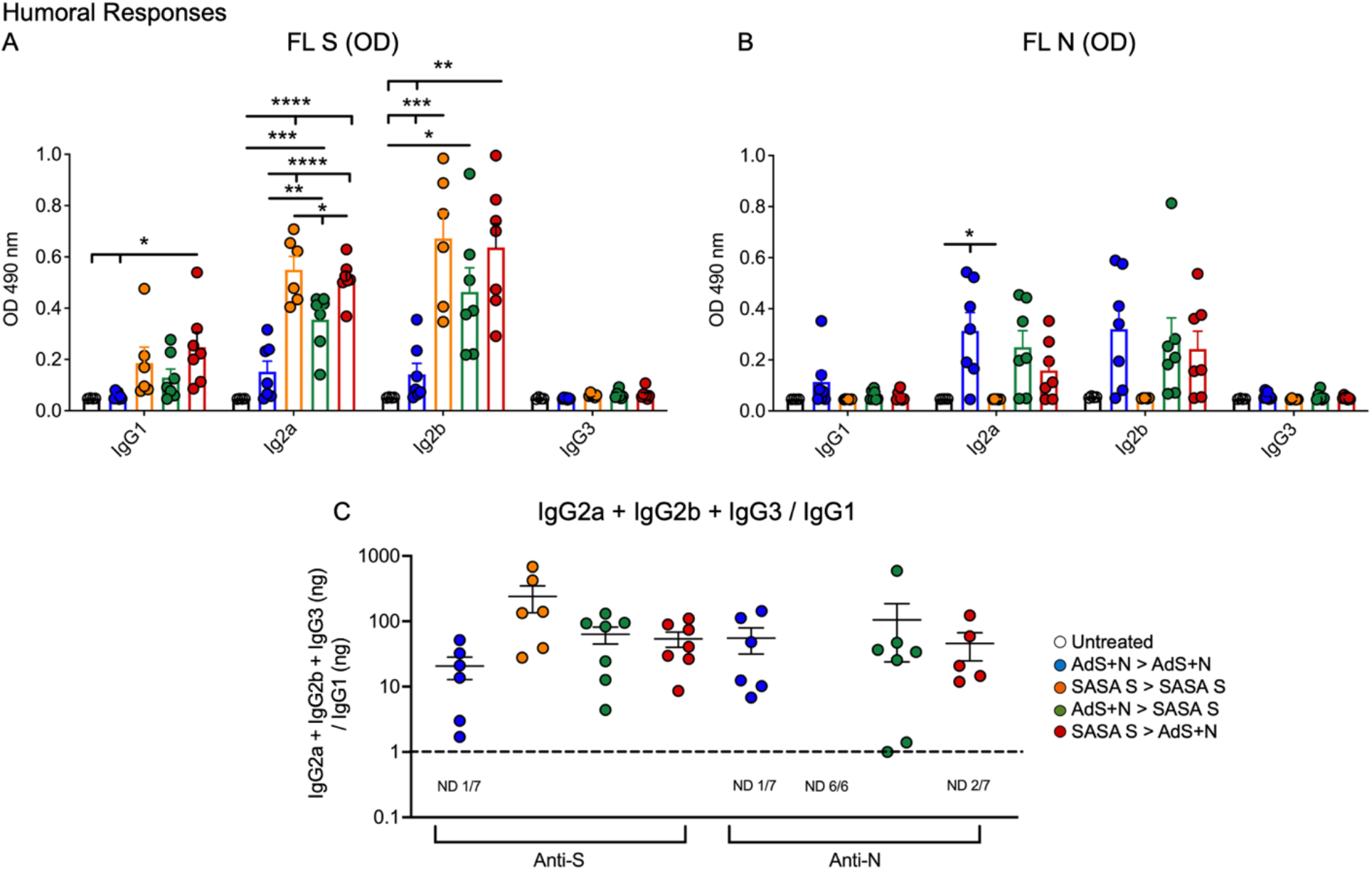
Anti-full length (FL) spike wild type (Swt) and -nucleocapsid (N) IgG antibody levels in sera show T helper cell 1 (Th1) bias. (A) Levels of anti-FL Swt and (B) anti-N IgG1, IgG2a, IgG2b and IgG3 subtypes represented by OD at 490 nm from ELISA of sera are shown. Statistical analyses were performed using one-way ANOVA and Tukey’s post-hoc comparison of all groups to all other groups with the exception of comparison to the SASA S > SASA S group that did not receive an N antigen for anti-N IgG; where *p ≤ 0.05, **p < 0.01, ***p < 0.001, and ****p<0.0001. (C) The IgG2a+IgG2b+IgG3/IgG1 ratio calculated using the ng equivalents for each is shown with a dashed line at 1. Values > 1 reflect Th1 bias. The number (n) of animals in which the ratio was not determined due to very low antibody levels is shown below the x-axis for each group. The homologous SASA S group did not receive an N antigen. Data graphed as the mean and SEM. The legend in C applies to all figure panels.

### Humoral Responses Against Wildtype and Delta S1 were Similar in All SASA S Groups

To assess serum antibody production specific for delta B.1.617.2 variant as compared to wild type S, an ELISAs were performed using either the wt or B.1.617.2 sequence S1 domain of S, which contains the RBD.

Vaccine regimens that included the SASA S vaccine elicited the highest anti-S1(wt) and S1(Delta) responses, as represented by the Ab50, μg IgG/mL, and endpoint titers (**Fig. 4A, B, and C**, respectively). Four of seven AdS+N homologous vaccinated mice had serum S-binding IgG levels that were below the level of detection. Overall, the mean antibody titers for SASA S homologous and SASA S > AdS+N groups were highest. For Ab50 and μg IgG/mL (**Fig. A and B**), statistical comparison of the AdS+N group to other groups was not performed because of the presence of values below the LOD in the AdS+N group. As measured by endpoint titer (**Fig. 4C**), anti-S1(wt) IgG responses were significantly improved when vaccination included an administration of SASA S, when compared with AdS+N homologous vaccination. Likewise, anti-S1(delta) IgG responses were significantly improved in animals with either SASA S homologous or SASA S > AdS+N vaccination, when compared with AdS+N homologous vaccination. These statistical analyses were performed by assigning reciprocal dilution endpoint titer values of 200 to those animals with IgG titers below the LOD.

**Fig. 4.**
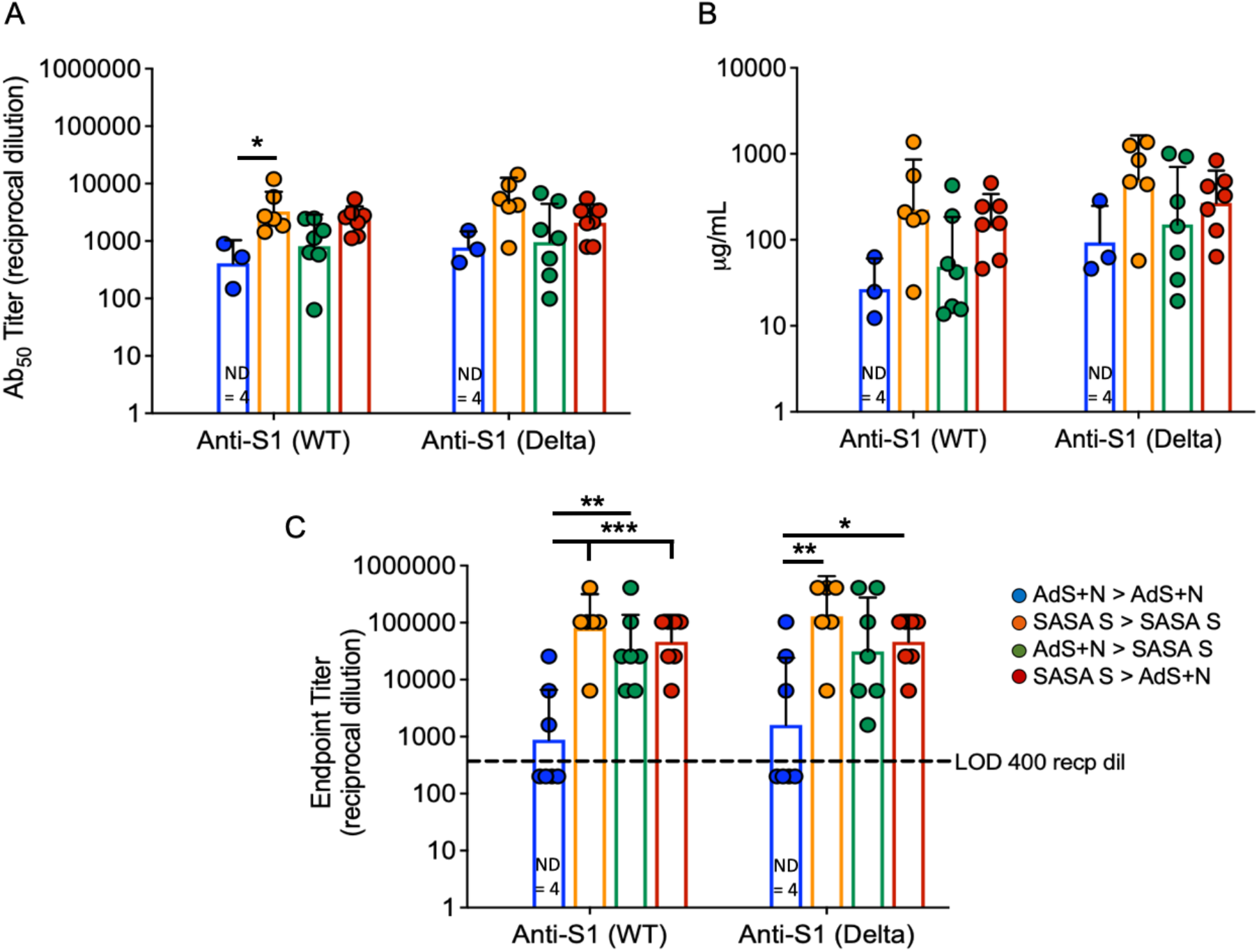
Wildtype and B.1.617.2 ‘Delta’ S1-specific IgG endpoint titers. Levels of anti-S1(wt) and – Delta S1 IgG are shown by (A) Ab50 reciprocal dilution, (B) μg/mL sera, and (C) endpoint titer reciprocal dilution. Values were below the level of detection in 4 of 7 AdS+N homologous group mice. Statistical analyses were performed on log-normalized data using one-way ANOVA and Tukey’s post-hoc comparison of all groups for anti-S1 (WT) or -S1 (Delta) where * p ≤ .05, **p < .01, and ***p < .001; sera without detectable levels of anti-S1 IgG were assigned a value of 200, one-half the Limit of Detection (LOD) of 400. Data graphed as the geometric mean and 95% CI. The legend in C applies to all figure panels.

### An AdS+N Boost after SASA S Prime Vaccination Enhances CD4+ and CD8+ T cell Responses

Significantly higher percentages of CD4+ T-cells secreting IFN-γ alone, IFN-γ and tumor necrosis factor-a (TNF-a), or IFN-γ, TNF-a, and interleukin-2 (IL-2) as detected by intracellular cytokine staining (ICS) in response to S(wt) peptides were detected in SASA S > AdS+N group mice as compared to both the AdS+N or SASA S homologous groups (**Fig. 5A, C, and D**). In addition, the mean percentages were significantly higher than for the AdS+N > SASA S group.

**Fig. 5.**
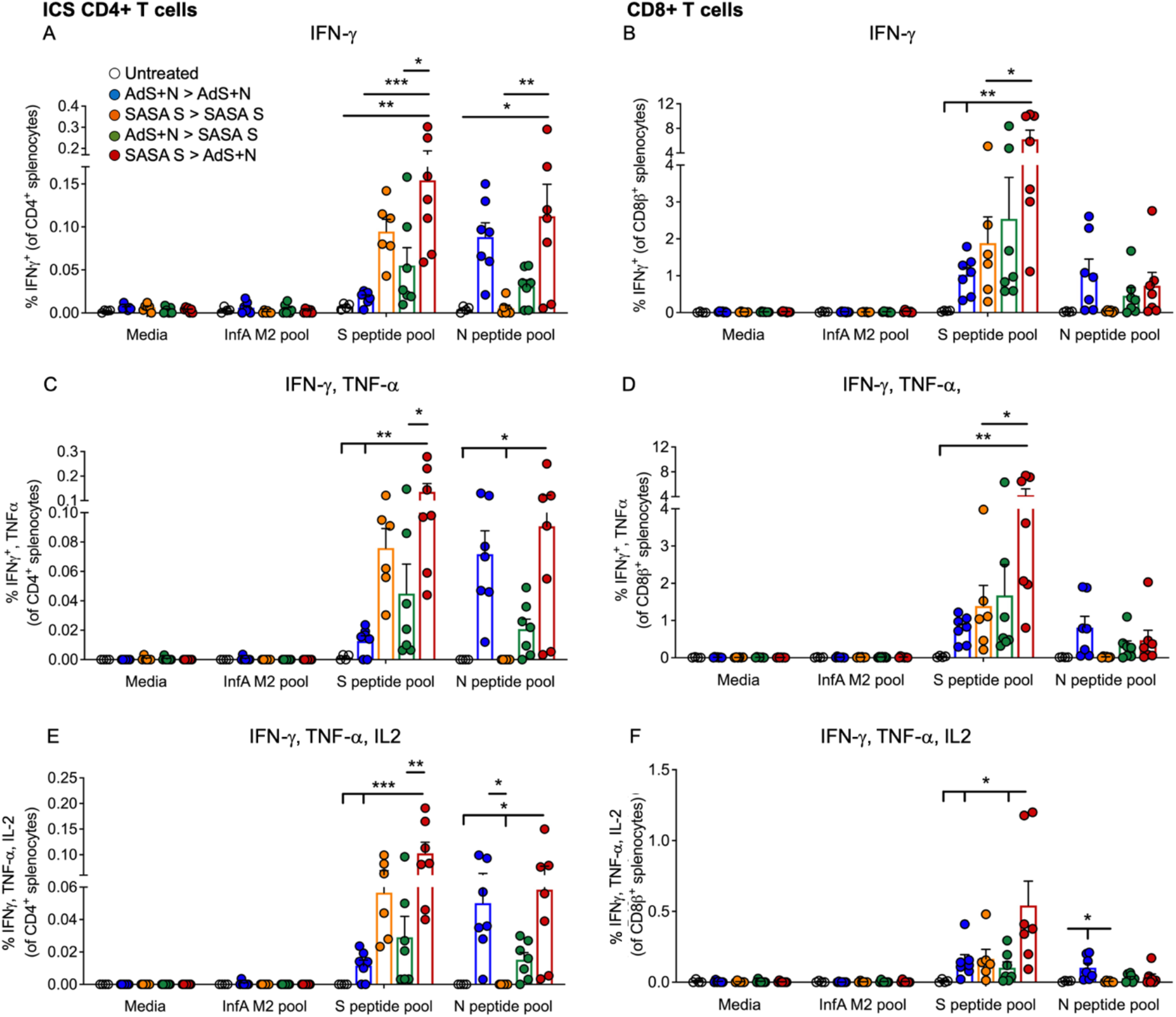
CD4+ and CD8+ T cell Intracellular cytokine staining (ICS) in response to S(wt) and N peptides. (A, B) ICS for interferon-γ (IFN-γ), (C, D) IFN-γ and tumor necrosis factor-a (TNF-a), and (E, F) IFN-γ, TNF-a and interleukin-2 (IL-2) are shown for CD4+ and CD8+ T cells, respectively. Statistical analyses performed using one-way ANOVA and Tukey’s post-hoc comparison of all groups to all other groups with the exception of comparison to the SASA S > SASA S group that did not receive an N antigen (comparisons to UnTx > UnTx are shown); where *p ≤ 0.05, **p < 0.01, and ***p < 0.001. Data graphed as the mean and SEM. The legend in A applies to all figure panels.

The enhancement of cytokine production by AdS+N boost of an SASA S prime was even more pronounced for CD8+ T cells (**Fig. 5B, D, and F**). Cytokine production was significantly higher in the SASA S > AdS+N group compared to both the homologous vaccination groups as well as the AdS+N > SASA S group.

As expected, only T cells from mice receiving vaccination regimens that included delivery of the N antigen by the AdS+N vaccine produced cytokines in response to N peptide stimulation. Mean responses of both CD4+ and CD8+ T cells to N peptides were similar for groups receiving AdS+N as a boost, either as part of homologous or heterologous vaccination (**Figure 5A-F**).

### CD4+ and CD8+ T-Cell Production of IFN-χ was Similar in Response to Either S(wt) or S(Delta) Peptides

CD4+ and CD8+ T cells show similar levels of IFN-γ production by ICS in response to either S(wt) or S(Delta) sequence peptides (**Fig. 6A and B**, respectively). Patterns of CD4+ and CD8+ T-cell stimulation by S protein peptides between the vaccination regimens were also similar between the S(wt) and S(Delta) peptides. Compared to the untreated control, the increase in IFN-g production was again the highest for the SASA S > AdS+N group for both CD4+ and CD8+ T cells, in response to either S(wt) or S(Delta) peptides.

**Fig. 6.**
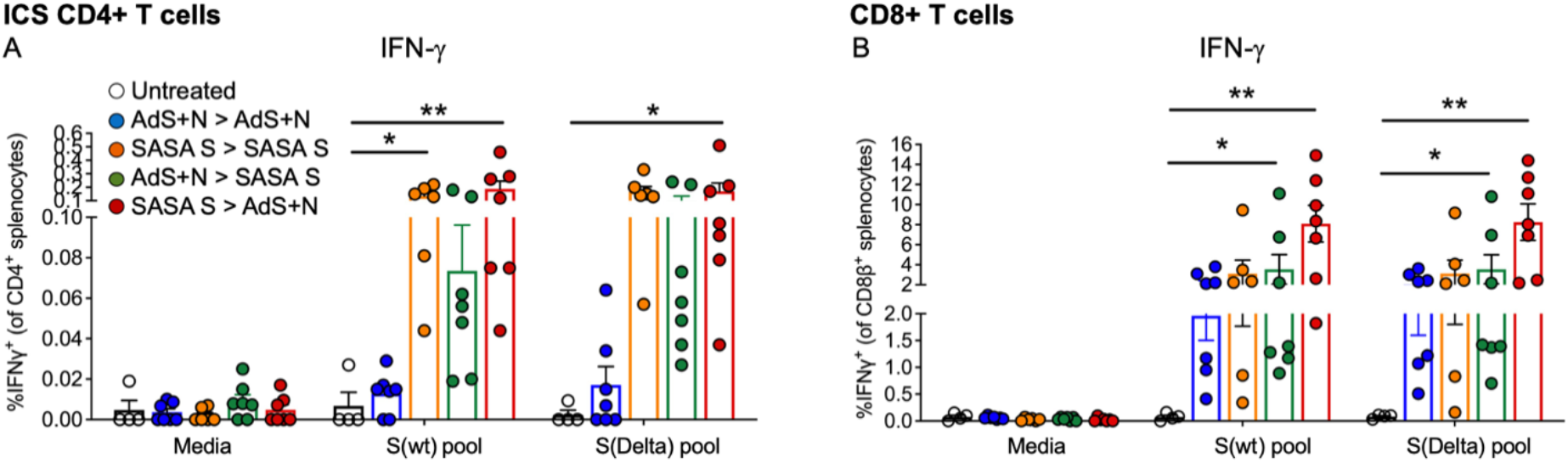
CD4+ and CD8+ T-cell responses to S(wt) and S(Delta) peptides are similar. Both CD4+ (A) and CD8+ (B) T cells show similar levels of interferon-γ (IFN-γ) production in ICS in response to either S(wt) or S(Delta) sequence peptides. For both T-cell types, the greatest responses were seen with SASA S > AdS+N vaccination. Statistical analyses performed using one-way ANOVA and Tukey’s post-hoc comparison of all groups to all other groups; where *p ≤ 0.05 and **p < 0.01. Data graphed as the mean and SEM. The legend in A applies to all figure panels.

### Numbers of IFN-γ-Secreting Splenocytes were the Highest from Mice Receiving SASA S > AdS+N Heterologous Vaccination

As shown in **Figure 7A**, ELISpot detection of cytokine secreting cells in response to S peptide stimulation revealed that animals receiving heterologous SASA S > AdS+ N vaccination developed significantly higher levels of S peptide-reactive IFN-γ-secreting T cells than all other groups except the SASA S homologous group (which had a lower mean, though statistical significance was not achieved). Numbers of IFN-γ-secreting T cells in response to the N peptide pool were similar for AdS+N homologous and SASA S > AdS+N groups. T cells from SASA S > SASA S group animals did not secrete IFN-γ in response to the N peptide pool, as expected, because the SASA S vaccine does not deliver the N antigen. While the difference was not significant due to individual variation, the mean number of N-reactive stimulated cells secreting IFN-γ due to AdS+N > SASA S vaccination was lower than the other groups receiving a vaccine with N. There was some skew seen for data in Fig. 7A, with values for S WT/N of untreated = 2.0/0.0, AdS+N > AdS+N = 1.27/0.27, SASA S > SASA S = −0.53/2.45, AdS+N > SASA S = 1.89/1.4, and SASA S > AdS+N = 0.35/-0118. We note these are outbred mice with variance in MHC class and variable T-cell data is not unexpected.

Induction of interleukin-4 (IL-4) secreting T cells was low for all animals in all groups (**Fig. 7B**), therefore the IFN-γ/IL-4 ratio was above 1 for all animals for which the ratio could be calculated (**Fig. 7C**), reflecting the Th1-bias of all T-cell responses.

**Fig. 7.**
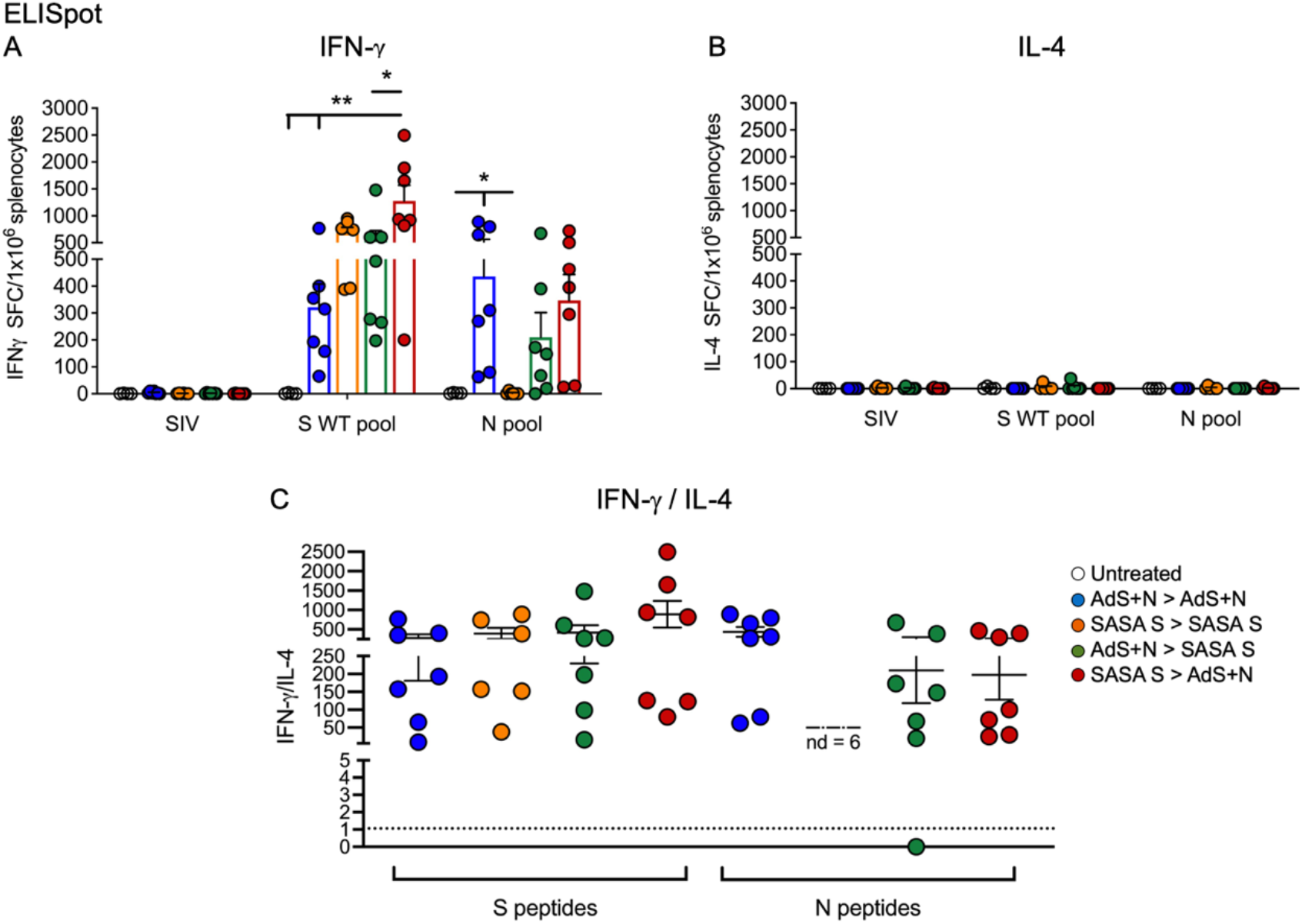
Heterologous vaccination increases T-cell cytokine secretion in ELISpot. (A) Numbers of interferon-γ (IFN-γ) and (B) interleukin-4 (IL-4) secreting T cells in response to S WT and N peptides pools. (C) The IFN-γ/IL-4 ratio; value of 1 indicated by dashed line. The ratio was not determined (ND) for animals with very low IL-4 secretion. Statistical analyses performed using one-way ANOVA and Tukey’s post-hoc comparison of all groups to all other groups where *p ≤ 0.05 and **p < 0.01. Data graphed as the mean and SEM. The legend in C applies to all figure panels.

### Sera from Mice Receiving the SASA S Vaccine Neutralize SARS-CoV-2 Wuhan, Delta, Beta and Omicron Pseudoviruses

As represented in **Figure 8A**, sera from SASA S homologous and SASA S > AdS+N heterologous group mice showed the highest neutralization capability (with AdS+N > SASA S being slightly lower) against the four SARS-CoV-2 lentiviral pseudoviruses: Wuhan (D614G), Beta (B.1.351), Delta (B.1.617.2), and Omicron (B.1.1.529) variants. Neutralizing antibody titers were significantly higher than for sera from untreated and AdS+N > AdS+N group mice.

**Fig. 8.**
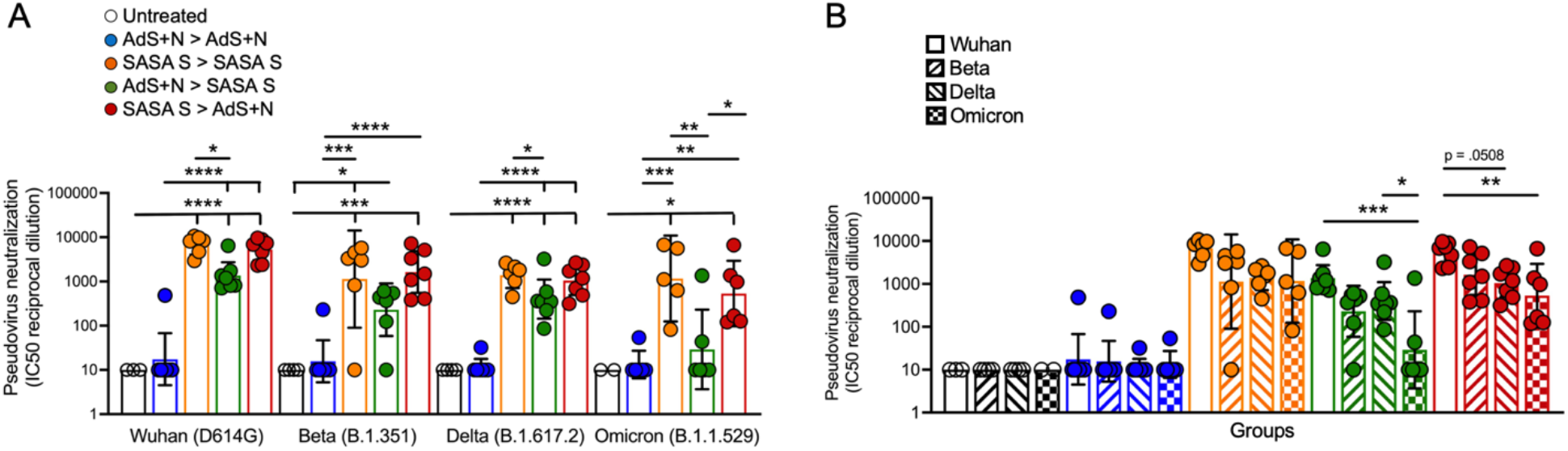
Sera from SASA S > AdS+N heterologously vaccinated mice neutralizes Wuhan, Delta, Beta, and Omicron strain SARS-CoV-2 pseudovirus. (A) IC50 reciprocal dilution for pseudovirus neutralization grouped by pseudovirus variant assay is shown. Statistical differences are shown for comparison of each vaccinated group for a specific variant (not between variants). (B) IC50 reciprocal dilution for neutralization of all strains/variant tested compared for each group is shown. The color code legend in (A) applies also to (B). Statistical comparison of IC50 values for untreated and AdS+N homologous group mice with values < the LOD was not performed. Statistical analyses were performed on log-normalized data using one-way ANOVA and Tukey’s post-hoc comparison where *p ≤ .05, **p < .01, ***p < .001, and ****p < .0001. Data graphed as the geometric mean and 95% CI.

Comparison of SARS-CoV-2 variant neutralizing antibody titers between groups (**Fig. 8B**) demonstrate that sera from SASA S homologous, AdS+N > SASA S and SASA S > AdS+N heterologous vaccinated mice all have high Wuhan-strain neutralization capacity. Sera from SASA S homologous vaccinated mice was capable of neutralizing all 4 strains similarly, but sera from both heterologously vaccinated groups showed a greater capability to neutralize the Wuhan strain versus the Omicron strain.

## DISCUSSION

Both CD4+ and CD8+ T-cell responses were enhanced by heterologous vaccination, with CD4+ interferon-γ (IFN-γ) production in response to both S(wt) and S(Delta) peptides highest with the SASA S prime > AdS+N boost combination as compared to all other groups. Notably, CD4+ and CD8+ T cells were equally responsive to S(wt) and S(Delta) peptides. Tesponses of T cells from SASA S > AdS+N to S(Delta) were also the highest of the groups.

Findings were similar by T cells ELISpot analyses, which again revealed the SASA S > AdS+N combination resulted in significantly higher IFN-γ secretion by T cells in response to both S peptides pools than all other groups.

We further demonstrate that all vaccine combinations that included the SASA S vaccine elicited the greatest anti-full length (FL) S wild type (wt), anti-S1(wt) and – importantly-anti-Delta variant (B.1.617.2) S1 IgG responses. Vaccine regimens that included the SASA S vaccine also generated sera that strongly neutralized Wuhan, Beta, Delta, and Omicron variant pseudoviruses.

As expected, anti-N IgG antibodies and T-cell responses to N peptides were seen only for vaccine combinations that delivered the N antigen and were very similar among groups receiving the AdS+N vaccine in any order.

The immune responses observed in the present study support our hypothesis that heterologous vaccination provides an opportunity for increased humoral and cell-mediated responses to vaccination. These results and hypothesis are consistent with recently-published data supporting enhanced antibody responses in patients who received heterologous vaccination with the currently-FDA authorized COVID-19 vaccines [36]. Delivery of the SASA S vaccine elicited higher anti-S IgG than the AdS+N vaccine, but the AdS+N vaccine provided the N antigen that broadened humoral responses and thus the potential to enhance protection against future SARS-CoV-2 variants of concern that could emerge. We note that mean anti-N IgG responses, while not statistically different among groups that received the N antigen (not SASA S homologous) were, as predicted, highest with homologous AdS+N vaccination.

Perhaps the most striking finding in the present study are the enhanced responses of S-specific CD4+ and CD8+ T-cells in SASA S > AdS+N group mice, an effect that was most pronounced for CD8+ T cells. The similarity of responses of CD4+ and CD8+ T cells to either S(wt) or S(Delta) suggest this vaccine regimen has a high probability of conferring T-cell mediated protection against the highly transmissible Delta variant – and by extension other emerging variants – in addition to humoral protection.

Immune responses elicited by SASA S > AdS+N vaccination were consistently the highest of the groups tested (although not always significantly so) and we hypothesize that because the SASA S vaccines elicits the greatest humoral response to S when given in any order -possibly reaching the upper detection limit for our ELISA – it enhances CD4+ T-cell activation as these are closely related to humoral/B cell responses. Therefore, CD4+ T-cell activation might be expected to be higher after a boost if there are stronger pre-existing, prime-induced B cell responses, that is, when SASA S is the prime. Adenovirus vectors such as that used for the AdS+N vaccine are good at eliciting CD8+ T-cell responses, which we posit explains why CD8+ T-cell responses are only slightly lower for homologous AdS+N vaccination (despite antibody and CD4+ T-cell responses being lower), as compared to heterologous vaccination. CD8+ T-cell responses likely also benefit from more robust pre-existing CD4+ T-cell and B cell responses, a condition that exists most prominently when the SASA S vaccine is given as the prime.

Effectively, enhanced CD4+-specific T-helper responses seen with SASA S prime dosing might have provided conditions for the enhanced CD8+ specific response upon AdS+N boost. The confirmation of this hypothesis awaits further investigation.

Importantly, all of the vaccination regimens that included the SASA S vaccine neutralized SARS-CoV-2 variant pseudoviruses – including the highly transmissible Omicron variant – reflecting the strength of humoral responses to the SASA S vaccine. We observed that the geometric mean IC50s for reciprocal dilutions of sera from mice receiving the heterologous SASA S > AdS+N regimen were consistently higher than those for AdS+N > SASA S, and speculate that the SASA S as a prime triggers greater B cell priming and development (as compared to AdS+N as the prime) which then results in enhanced recall when the AdS+N boost is delivered.

The lower capability of sera from AdS+N homologously vaccinated mice does not necessarily indicate that the predominantly T-cell inducing AdS+N vaccine would not be effective in protecting against SARS-CoV-2 challenge; in fact, we have previously reported that homologous AdS+N prime-boost vaccination of non-human primates confers protection against viral challenge [7]. In the *in vivo* viral challenge testing paradigm, cell-mediated immunity – not accessed in the pseudovirus assay that tests sera – conferred by AdS+N vaccination likely plays a key role in protection, as has been reported for natural infection of patients [37–40].

The findings here, including cross-reactive humoral and T-cell responses to S Delta and, particularly for regimens including SASA S, neutralization of Wuhan, Beta, Delta, and Omicron pseudovirus, support ongoing studies of heterologous vaccination with the SASA S and AdS+N vaccines. Further testing in pre-clinical models of SARS-CoV-2 challenge and clinical trials should be conducted to assess the capability of this vaccine regimen to provide increased protection against COVID-19 and SARS-CoV-2 variants by combining the ability of SASA S to elicit vigorous humoral responses with AdS+N’s second, highly antigenic N antigen and T-cell response enhancement.

## ACKNOWLEDGEMENTS

We would like to thank Jesse Bloom (Fred Hutchinson Cancer Research Center) and Thomas Peacock (Imperial College London) for sharing the SARS-CoV-2 spike protein plasmids used for pseudovirus production.

## DATA AVAILABILITY STATEMENT

The datasets presented in this study can be found in online repositories. The names of the repository/repositories and accession number(s) can be found in the article and include Rice et al. 2021 [41].

## ETHICS STATEMENT

The animal study was reviewed and approved by the Institutional Animal Care and Use Committee (IACUC) at Omeros, Inc. (Seattle, WA, USA).

## AUTHOR CONTRIBUTIONS

AR and MV contributed to the study design, co-wrote the manuscript and, with KD and SM, performed the in vivo studies and co-analyzed data; EV is co-inventor of the RNA technology used for the SASA S saRNA vaccine, contributed to design of the study, co-analyzed data and data interpretation, and edited the manuscript; SB performed the pseudovirus neutralization assay, with the assistance of PB and SR, and edited the manuscript; LZ, CAO, ST, and BM contributed to the design, production, and testing of the AdS+N vaccine; EG co-designed the AdS+N vaccine vector; JTS contributed to the study design and provided expert immunological/biological insight for interpretation of data; PS analyzed data, generated figures and tables, and wrote the manuscript; CC contributed to the study design and data analysis, and edited the manuscript; PS-S codesigned and developed the AdS+N vaccine, co-conceptualized the study, reviewed all data, and edited the manuscript. All authors contributed to the article and approved the submitted version.

## FUNDING

The original development of the SASA S vaccine was funded by the Infectious Disease Research Institute (IDRI). The development of the hAd5 S-Fusion+N-ETSD (AdS+N) vaccine and the. present study was funded by ImmunityBio, Inc.

## DISCLOSURES

All authors with an ImmunityBio, Inc., affiliation contribution to the design, production or testing of the AdS+N vaccine that may become a commercial product. Emily Voigt is an inventor on a patent related to the RNA vaccine technology.

## Notes

### Competing Interest Statement

The hAd5S+N and SASA S vaccines are under development by ImmunityBio, Inc.

### Summary of Updates

Data for sera neutralization of omicron pseudovirus had been added, some statistics were re-performed on log-normalized data in those instances where statistics were performed on data values graphed on a log scale.

